# An amplitude code increases the efficiency of information transmission across a visual synapse

**DOI:** 10.1101/328682

**Authors:** Ben James, Léa Darnet, José Moya-Díaz, Sofie-Helene Seibel, Leon Lagnado

## Abstract

**Most neurons in the brain transmit information digitally using sequences of spikes that trigger release of synaptic vesicles of fixed size. The first stages of vision and hearing are distinct in operating with analogue signals, but it is unclear how these are recoded for synaptic transmission. By imaging the release of glutamate in live zebrafish, we demonstrate how ribbon synapses of retinal bipolar cells transmit analogue visual signals by changes in both the rate and amplitude of synaptic events. Higher contrasts released glutamate packets composed of more vesicles and coding by amplitude often continued after rate coding had saturated. Glutamate packets equivalent to five vesicles transmitted four times as many bits of information per vesicle compared to independent release events. By discretizing analogue signals into sequences of numbers ranging up to eleven, ribbon synapses increase the dynamic range, temporal precision and efficiency with which visual information is transmitted.**

---

The spike code of neurons has been studied in detail^1,2^ but much less is known about the vesicle code transmitting information across the synapse^3^. Since the work of Katz^4-6^, the general view has been that changes in presynaptic potential are communicated by modulating the mean rate of a Poisson process in which vesicles at the active zone are triggered to fuse independently^7-9^. At most synapses, the voltage signal controlling this process is digital, arriving in the form of an all-or-nothing spike, and the output is also digital, in the form of a vesicle releasing a fixed packet of neurotransmitter^2^. But in the first stages of vision and hearing the neural signal is in analogue form - continuous changes in membrane potential that are graded with the strength of the stimulus ^10^. How are these sensory signals recoded for transmission across a synapse?

Synapses driven by analogue signals are distinguished from those driven digitally by the presence of a specialized structure, the ribbon, that holds many tens of vesicles just behind the active zone^11,12^. It has generally been assumed that these ribbon synapses carry out the key step of analogue-to-digital conversion when the graded voltage signal is represented as changes in the rate of events releasing just one quantum of neurotransmitter^7,8,13-15^. A number of *in vitro* studies have, however, shown that ribbon synapses are also capable of modulating the size of the packet of neurotransmitter through a process termed coordinated multivesicular release (CMVR)^15-20^. This observation is intriguing because it suggests that the output of a ribbon synapse may not be a single “symbol”, but a number of symbols varying in amplitude^21^. The functional role of CMVR has, however, been unclear because it has not been observed in response to light or sound.

To understand whether the coordinated release of vesicles contributes to the transmission of visual information we used the fluorescent glutamate reporter iGluSnFR^22^ to image the output of ribbon synapses in the retina of zebrafish. Optimizing the signal-to-noise ratio allowed us to count vesicles released from individual active zones *in vivo* and investigate how they are used to transmit the visual signal while it is still in analogue form. We show that ribbon synapses of bipolar cells employ a coding strategy that is a hybrid of the known rate code and of changes in the amplitude of synaptic events, which we term the amplitude code. We describe how amplitude coding confers several advantages, including the ability to signal contrast beyond the range where the rate code has saturated, improving the temporal precision of transmission and increasing the efficiency of communication by transmitting more bits of information per vesicle.

## Results

### Optical detection of co-ordinated multivesicular release *in vivo*

All visual signals flow through bipolar cells because these are the only neurons that connect the photoreceptors to ganglion cells the inner retina (Fig. 1a). To investigate how these signals are represented as they cross the synapse, expression of iGluSnFR was driven in bipolar cells using the *ribeye* promoter^14^. A periodic stimulus (5 Hz) triggered a train of glutamate transients which were sampled by performing line scans through the terminal at 1 kHz (Fig 1b and c; Video S2). These transients were generated by the electrical signal arriving from the cell body rather than glutamate spillover from neighbouring synapses because they were destroyed by ablating the soma of the cell (Fig. S8). Indeed, closer analysis of the intensity profile along a line demonstrated that iGluSnFR signals were constrained to spatial scales of ∼1-2 mm making it possible to distinguish fusion events at neighbouring active zones by spatial demixing (Fig. 1c and Supplementary Figure 2). iGluSnFR signals at adjacent active zones did not always coincide in time, reflecting the stochastic nature of vesicle release (Fig. 1c) and, most notably, the size of glutamate transients varied widely (Fig. 1c and d).

**Figure 1.**
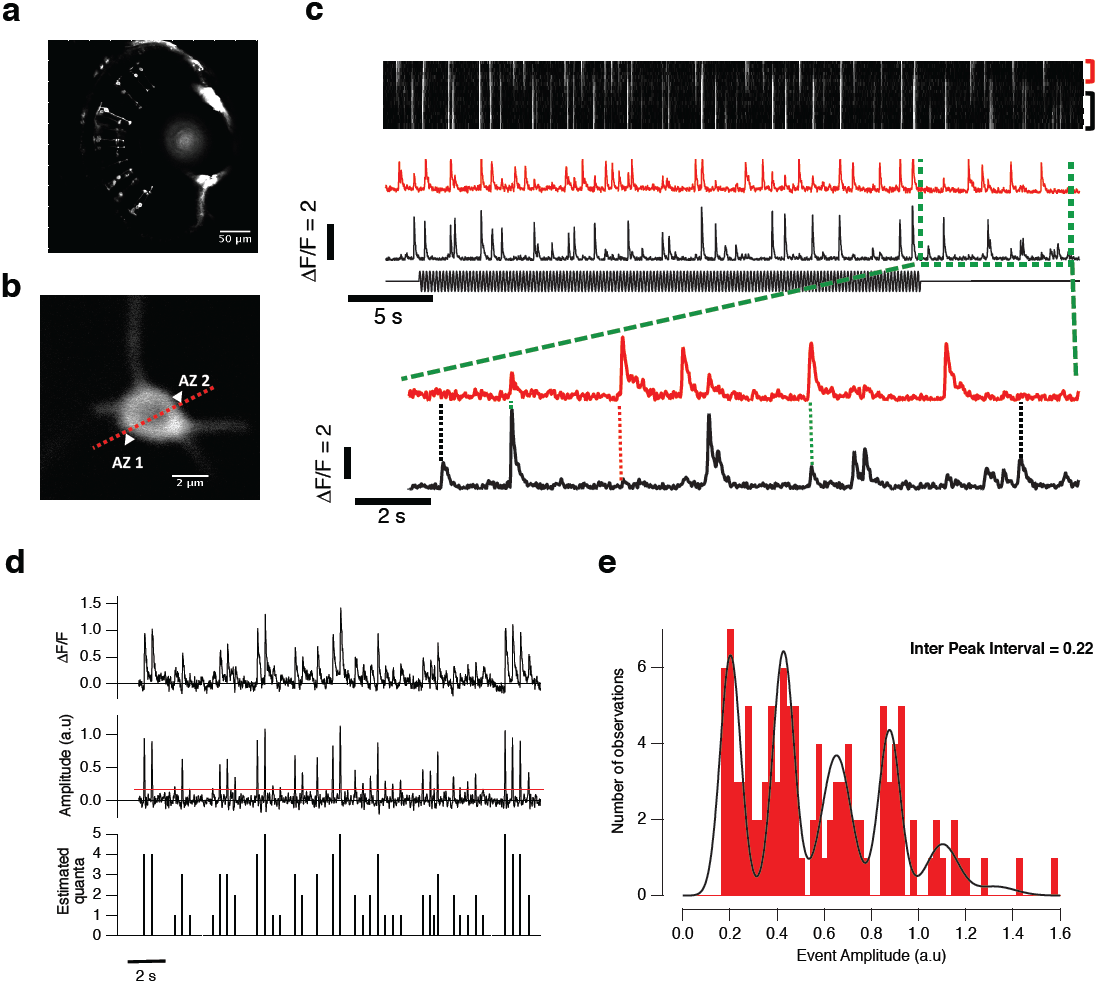
Quantized release of glutamate imaged at individual active zones. **a)** Multiphoton section through the eye of a zebrafish larva expessing iGluSnFr in bipolar cells at 7dpf. **b)** Linescan through a single terminal. Arrowheads shows the position of two active zones. **c)**. The kymograph (top) shows the intensity profile along the line in B as a function of time. The red and black traces (middle) show the time-course of the iGluSnFR signal over the two active zones marked to the right of the kymograph and the stimulus is shown immediately below. After 2 s there was a switch from constant illumination to full field modulation at 20% contrast, 5 Hz. A section of the record (green box) is expanded Note variations in the amplitude of glutamate transients. Sometimes both active zones release glutamate (green dashed line) while on other occasions only AZ 1 (black dash) or AZ 2 (red dash). **d)** Raw (top) and deconvolved (middle) traces from a synapse stimulated at 50% contrast. Transients in the deconvolved trace were considered for analysis if they crossed a threshold (red line). The bottom panel shows the estimation of the number of quanta per event, based on the quantal size estimated in e. **e)** Amplitude histogram extracted from the deconvolved trace in **d**. The red line is a fit of six Gaussians, constrained to have equal distance between each peak but with all other parameters free. The peaks are separated by an inter-peak distance of ∼0.22, which is the amplitude of the event equivalent to a single vesicle.

To improve estimates of the timing and amplitude of glutamatergic events, a Wiener filter was applied^9^ in which the raw iGluSnFR trace (Fig. 1d, top) was deconvolved (middle) using a temporal kernel estimated by averaging a number of isolated events (Supplementary Figures 3 and 4; described in detail in the Supplementary Methods). Histograms of event amplitudes measured from deconvolved traces displayed several peaks at equal intervals indicating that transients of different sizes were composed of multiples of an underlying quantal event (Fig. 1e). These quanta are very likely to correspond to synaptic vesicles given the electrophysiological evidence that ribbon synapses in auditory hair cells and the retina can sometimes coordinate the fusion of two or more vesicles rather than releasing them all independently^16,17,19^. Based on amplitude histograms allowing estimates of the quantal signal, we partitioned iGluSnFR signals of different sizes into numbers of quanta using a maximum likelihood estimate, as shown in the lower panel of Fig. 1d and described in Supplementary Methods.

Two further observations indicated that larger glutamate events involved the synchronized release of vesicles rather than coincidental and independent fusion occurring within a narrow time window^16-19^. First, the time-courses of iGluSnFR signals containing multiple quanta were indistinguishable from those containing just one (Supplementary Figure 4). Second, the distribution of time intervals between events consistently deviated from the Poisson statistics expected for vesicles released independently with constant probability (Supplementary Figure 9). Coordinated release of two or more vesicles was observed in all of 186 active zones that we investigated in both the ON and OFF channels of the retina (74 ON and 112 OFF synapses).

### Reverse-correlation at an active zone: the transmitter-triggered average

The existence of a process that synchronizes the release of two or more vesicles has been recognized at ribbon synapses for some years, but it’s functional role has not been clear^16-19^. Particularly puzzling has been the observation that, although the frequency of CMVR in hair cells and bipolar cells is calcium-dependent, the amplitude of events is not^17,18^. At face value, this would suggest that CMVR does not encode the loudness of sound or contrast of a visual stimulus. These experiments did not, however, measure responses to a physiological stimulus but instead stimulated the synapse through a patch pipette^17,18^. Counting vesicles with iGluSnFR allowed us to make an *in vivo* investigation of how CMVR contributes to the encoding of visual stimuli.

We began investigating this question by adapting an approach that has been widely used to explore the information represented by spikes – calculation of the spike-triggered average (STA). The STA is calculated by reverse-correlating the spike train generated by a random (“white noise”) stimulus to the stimulus itself^2,23^ and provides an estimate of the neurons tuning in the form of its linear filter in time or in space and time. By reverse-correlating events measured using iGluSnFR in response to full-field noise, we instead constructed the “transmitter triggered average” (TTA) to describe the output of an active zone. This approach to understanding the function of a synapse was first proposed in 2004^3^, but has not been realized until now. A fundamental distinction between the STA and TTA was that the while the spike output of a neuron contains just one symbol, here we are dealing with a vesicle code involving a number of symbols (1, 2, 3 quanta etc). We therefore calculated the TTA separately for synaptic events composed of different numbers of vesicles (Fig. 2a).

The TTA revealed that the more quanta within an event the higher, on average, the temporal contrast driving it (Fig. 2b). The relationship between the number of quanta released per event (Q_e_) and the contrast in the temporal filter (C) could be described by a first-order saturation of the form C = C_max_ x Q_e_ /(Q_e_ + Q_1/2_), where C_max_ = 19% and Q_1/2_ = 2.4 vesicles (Fig. 2c). Thus, CMVR encodes one of the most fundamental properties of a visual stimulus – temporal contrast – in a simple and direct way.

**Figure 2.**
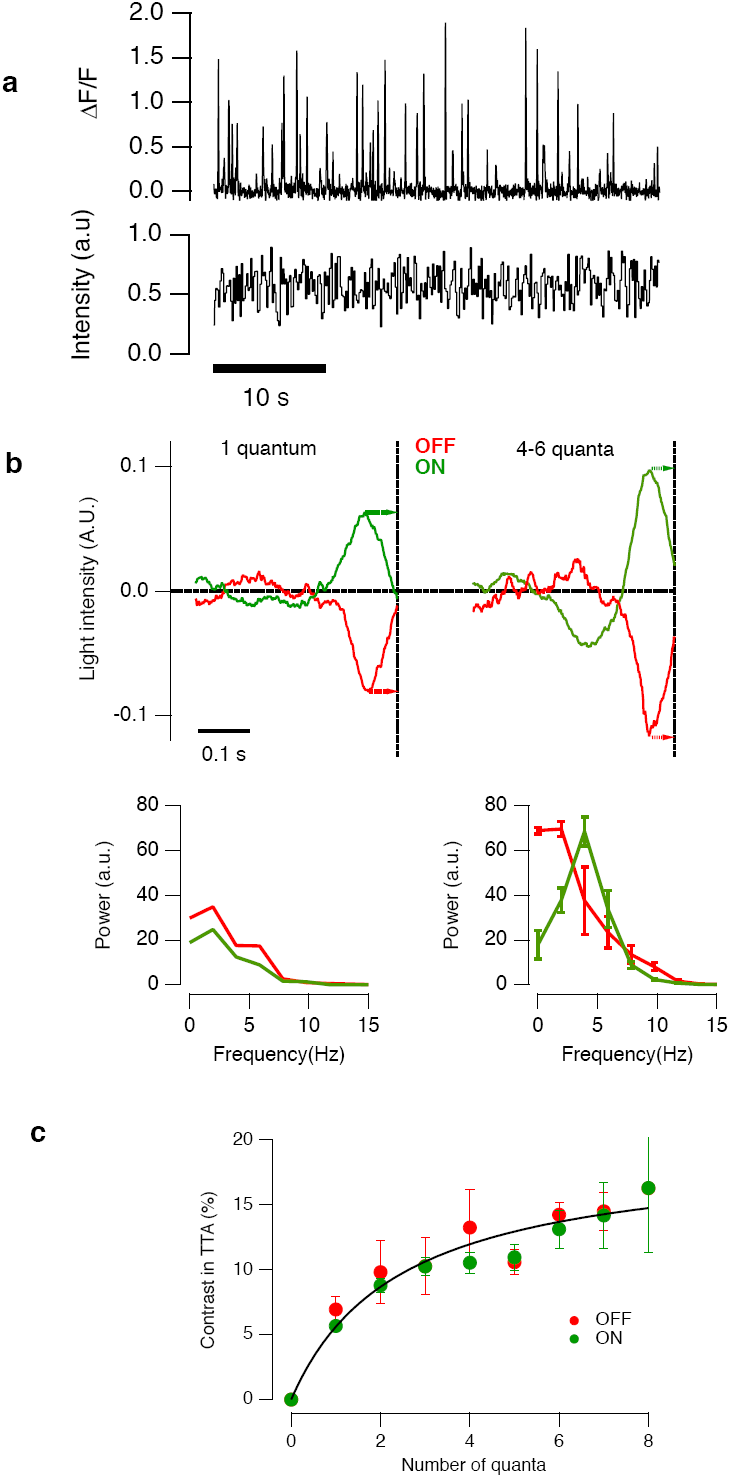
The transmitter-triggered average depends on the number of quanta in a release event. **a)** Example of iGluSnFR signals (top) elicited by a “white noise” stimulus (bottom). **b)** Upper traces: Linear filters extracted by reverse-correlation of responses composed of one quantum (left) and four to six quanta (right) from ON synapses (green, n = 9) and OFF synapses (red, n = 8). Multiquantal events encoded larger modulations in intensity. The plots below show the power spectra of the linear filters. Through the OFF channel, both uniquantal and multiquantal events were driven through low-pass filters, but in the ON channel multiquantal events were driven through a band-pass filter peaking at ∼4 Hz. **c)** Relation between the Michelson contrast represented in the TTA and the number of quanta released within the events used for reverse correlation in ON and OFF terminals. The relation could be described by a first-order saturation of the form C = C_max_.N/(N + N_1/2_), where C_max_ = 19% and N_1/2_ = 2.4. Bars show standard errors.

The TTA revealed a second distinction between events of different amplitude: uniquantal events were characterized by temporal filters that were monophasic through both ON and OFF channels, corresponding to a low-pass filter, but the TTA from ON synapses were biphasic corresponding to band-pass characteristics with peak transmission at ∼4Hz (Fig. 2b). In other words, a period of hyperpolarization before depolarization favoured larger synaptic events through the ON channel. Such biphasic antagonism in the time-domain is commonly observed in the temporal receptive fields of neurons early in the visual system ^24^, where it has been proposed to underly the suppression of redundant signals ^25^. Synchronizing the release of multiple vesicles is expected to amplify the output of a bandpass filter to enhance the signaling of change in the time-domain. Together, the results in Figs. 1 and 2 demonstrate that CMVR plays a fundamental role in transmitting the visual signal^16,17,19^.

### A hybrid rate and amplitude code

To investigate further how changes in the amplitude of synaptic events transmitted information about a visual stimulus we constructed contrast-response functions for individual active zones. Fig. 3a shows a protocol used to quickly assess this function, in which the contrast of a 5 Hz stimulus was increased in 10% steps. The first measure of response that we used was the total number of quanta per cycle of the stimulus (Q_c_), from which we estimated the contrast generating the half-maximal response, C_1/2_, as shown by the example in Fig. 3b. This value occurs at the steepest part of the contrast-response function, and so also defines the range in which the synapse signaled a change in contrast with the highest sensitivity^2^. A first inspection of these traces shows that an increase in contrast increases both the frequency and amplitude of glutamatergic events, as can also be seen in the records in Fig, 4a, Fig. 5a and Fig. 6a.

**Figure 3.**
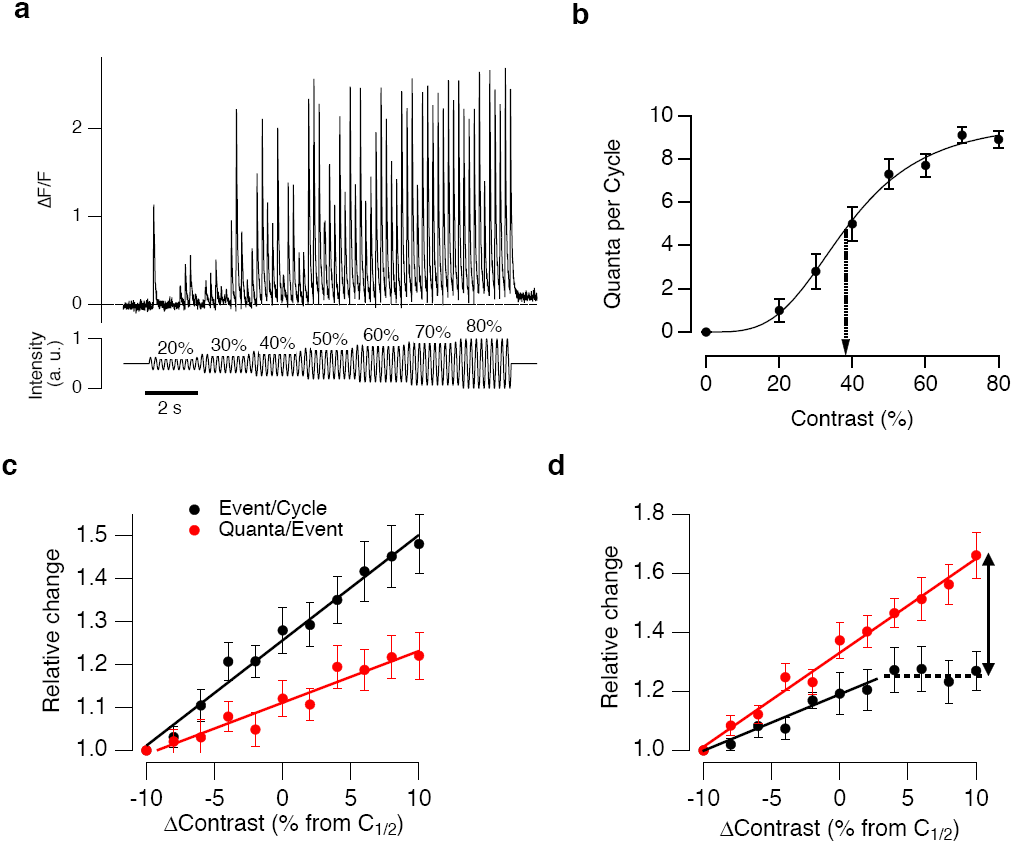
The relative contributions of coding by rate and amplitude. **a)** Example of the iGluSnFR signal (top) elicited by a full-field stimulus of increasing contrast delivered at 5 Hz (bottom). This protocol was used to quickly assess the half-point of the contrast-response function (C_1/2_). **b)** Contrast-response function extracted from the synapse in **a**, with response (R) quantified as the total number of quanta per cycle of the stimulus. The curve is a Hill equation of the form C = R_max_.(C^h^/C^h^ + C^h^), where R_max_ = 9.8 quanta per cycle (49 quanta s^−1^), h = 3.5 and C_1/2_ = 39% (dashed arrow). **c)** The relative change in synaptic activity around C_1/2_. The average number of events per cycle (E_c_, black) is compared with the average number of quanta per event (Q_e_, red). Stimuli were delivered in 2 s episodes with 2 s rest as shown in Fig. 6a. In these 38 synapses an increase in contrast caused E_c_ to rise more steeply than Q_e_. **d)** In the remaining 17 synapses, Q_e_ rose more steeply than E_c_, which then saturated (dashed line) such that further increases in contrast were signaled only by increases in the number of quanta per event (vertical arrow).

What are the relative contributions of the rate and amplitude codes? To assess how these two modes of signaling operated in parallel, we focused on the range where contrast sensitivity was highest, C_1/2_ ± 10%, using a second stimulus protocol in which different contrasts were applied in 2 s steps separated by 2 s intervals, as shown in Fig. 6a. Q_c_ was then factorized into two quantities: the number of events per cycle (E_c_, representing the contribution of the rate code) and the number of quanta per event (Q_e_, the amplitude code). The relative change in E_c_ and Q_e_ varied across synapses: in 38 of 55 active zones, an increase in contrast caused E_c_ to rise more steeply than Q_e_ (Fig. 3c) but in the remaining 17 the amplitude code was dominant and Q_e_ rose more rapidly than E_c_ (Fig. 3d). Synapses in which the amplitude of synaptic events was modulated more strongly displayed a second striking property: when the rate code saturated, further increases in contrast were represented *wholly* as increases in the size of the glutamate packets released (arrowed in Fig. 3d). CMVR therefore extends the range of contrasts that can be signaled beyond those allowed by the rate code alone.

One factor determining the shift towards release events composed of more quanta at higher contrasts was the ON/OFF identity of the bipolar cell. The change in the distribution of event amplitudes when the contrast was increased from 20% to 100% are shown in Fig. 4. At 20% contrast, Q_e_ was not significantly different in the ON and OFF channels (0.48 ± 0.19 and 0.58 ± 0.18; n= 18 and 33, respectively). But at 100% contrast, the average event amplitude in the OFF channel (5.07± 0.45 quanta) was significantly higher than the ON (2.89 ± 0.5; p<0.0001 students t-test). The distribution of event amplitudes at 100% contrast were also significantly different through ON and OFF channels (Chi-squared;p<0.0001). For instance, there was a cut-off above which larger events uniquely coded higher contrasts which was about six quanta and for the OFF channel and four quanta for the ON (Fig. 4b and c). The largest synaptic events in both ON and OFF bipolar cells were composed of about eleven vesicles (Fig. 4b). Thus, while most synapses in the brain signal with zeros and ones, the ribbon synapses of bipolar cells go up to eleven.

**Figure 4.**
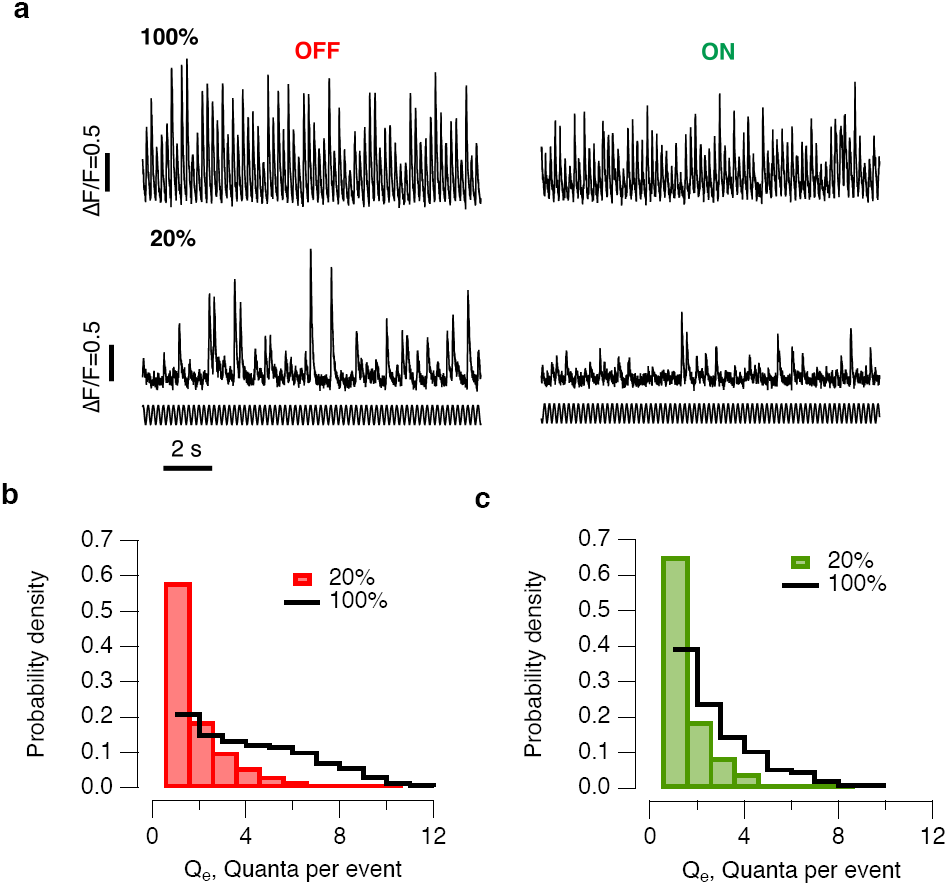
Changes in the distribution of event amplitudes. **a)** Representative traces of synaptic responses in ON and OFF terminals stimulated at 20% and 100% contrast (5 Hz). Note the increase in the frequency and average amplitude of events at higher contrast. **b)** Probability density of events composed of different numbers of quanta in OFF synapses (n = 33). Events composed of more than 6 quanta were only observed at contrasts greater than 20%. **c)** Distributions for ON synapses (n = 18). Events composed of more than 4 quanta were only observed at contrasts greater than 20%. The distribution at 100% contrast was significantly different to that observed in OFF synapses (Chi-squared; p<0.0001).

Ribbon synapses of bipolar cells therefore encode a visual stimulus through a hybrid strategy involving changes in both the rate and amplitude of synaptic events. These observations present a fundamentally distinct picture to that obtained by making electrophysiological measurements of CMVR from hair cell ribbon synapses *in vitro*, where the rate of events depended on the amplitude of modulation of the sinusoidal command voltage but the average event size did not^20^. It may therefore be that CMVR at the ribbon synapse of hair cells is not used to encode loudness, although further experiments in a more intact preparation would be required to confirm this (see Discussion).

### The amplitude code improves the temporal resolution of visual signals

Events of larger amplitude improved the temporal precision with which the visual signal was transmitted. The timing of uniquantal events relative to the phase of a 5 Hz stimulus is shown in Fig. 5a for contrasts of 20% and 100%. Uniquantal events displayed a standard deviation (“temporal jitter”) of 24 to 28 ms over a range of contrasts, while events composed of 7 or more quanta jittered by as little as 2.5 ms (Fig. 5b). The spike trains of retinal ganglion cells (RGCs) post-synaptic to bipolar cells can also encode visual signals of high contrast with millisecond precision^26^ and the stronger excitatory input from large glutamatergic events may be one mechanism by which this is achieved.

**Figure 5.**
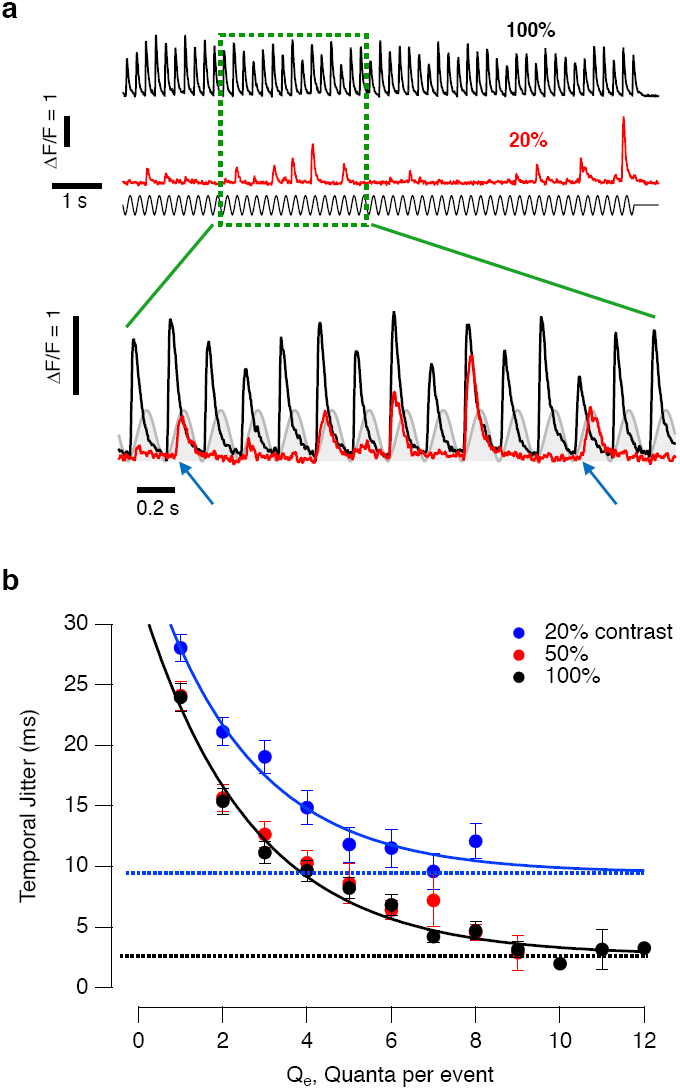
Multivesicular events increased the temporal precision of synaptic transmission. **a)** Top: iGluSnFR signals from a terminal stimulated at 100% and 20% contrast. Note the large variation in event amplitudes at lower contrast. Bottom: expanded time-scale showing responses at 20% and 100% contrast superimposed on the phase of the stimulus (grey). Events composed of fewer quanta are less synchronized than multiquantal events. Two examples of events occurring at phases different to the majority are shown by the blue arrows**. b)** Temporal jitter (s) as a function of the number of quanta per event (Qe) at contrasts of 20% (blue, n= 57 synapses), 50% (red, n= 61) and 100% (black, n= 63). Events composed of more quanta exhibit lower jitter at a given contrast and events of a given amplitude exhibit lower jitter at higher contrasts. The relationship between s and Q_e_ could be described as s = s_min_ + (s_max_-s_min_).(e^−Qe/k^). At 100% contrast, s_min_ = 2.7 ms (dashed black line), s_max_ = 30 ms and k = 2.6 quanta. At 20% contrast, s_min_ = 9.5 ms (dashed blue line), s_max_ = 28 ms and k = 2.4 quanta.

Stronger synchronization of larger synaptic events has also been observed by making electrophysiological measurements of the output from hair cell ribbon synapses driven by a sinusoidal command voltage^20^. This property can be quantified by a metric called the vector strength, which varies between zero (random occurrence of events) and one (perfect synchronization with the stimulus) ^27^. The vector strength (s) is related to the temporal jitter (r) by the relation s = (2(1-r))^1/2^/2Pf where f is the frequency. In the experiments shown in Fig. 5, the vector strength achieved a near perfect value of 0.98 at 100% contrast. In comparison, *in vitro* experiments in hair cells have measured multivesicular events to have vector strengths up to 0.6 at frequencies of 400 Hz^20^. These different degrees of synchronization might reflect the higher operating frequencies of auditory synapses compared to visual in combination with an inherent temporal limit in the processes triggering CMVR.

The degree to which synaptic events were consistent in time depended not only on event amplitude, but also the contrast of the stimulus eliciting the event (Fig. 5b). For instance, the largest packets of glutamate consistently observed at 20% contrast contained 8 quanta with a temporal jitter of 12.1 ± 1.4 ms, while 8 quantal events at 100% contrast jittered by 4.6 ± 0.6 ms. These observations run counter to electrophysiological measurements of CMVR made from hair cell ribbon synapses, where the amplitude of the changes in presynaptic potential had no effect on the size of the evoked events^20^. The properties of the amplitude code shown in Fig. 5 are, however, similar to the properties of the spike train in post-synaptic ganglion cells, which also become more temporally precise as the contrast is increased^26^. A shift to larger glutamatergic inputs of higher temporal precision may be one of the mechanisms that cause the spike trains of ganglion cells to become less variable as contrast is increased.

### Amplitude coding improves the efficiency of information transmission

The ionic mechanisms that generate neural signals and the synaptic processes that transmit them are a major energetic cost to the brain^28,29^. The need to transmit information in an energy-efficient manner has provided a unifying principle by which to understand the design of sensory circuits^15,30,31^. This framework provides, for instance, a teleological explanation for the use of analogue signaling to transmit early visual and auditory information: graded changes in membrane potential transmit information more efficiently than spikes^15^. The reason that analogue signals do not find more widespread use in the vertebrate nervous system is that they dissipate over relatively short distances, requiring a switch to digital signaling with regenerative spikes to relay information beyond the distance of one or two small neurons. The finding that ribbon synapses employ a hybrid rate and amplitude code to transmit analogue signals therefore led us to compare the amount of information contained within uniquantal and multiquantal events.

Using information theory^32^, we quantified the mutual information between a set of stimuli of varying contrasts (Fig. 6a) and release events containing different numbers of quanta (Fig. 6b). Vesicles released individually carried an average of 0.125 bits of information which is, as expected, significantly less than the 1-3.6 bits transmitted per spike in post-synaptic ganglion cells^26,33^. The amount of information contained within events of different size is shown in Fig. 6b. Larger events transmitted progressively more information because they were rarer^34^ (Fig. 4b and c) and driven preferentially by higher contrasts rather than occurring randomly (Fig. 2). The relation between the specific information (i, bits) and Q_e_, the number of vesicles comprising the event, could be described as a power function of the form i = i_o_ + A.Q_e_^x^, with i_o_ = 0.12 bits, A= 0.008, x = 2.8. This supralinear relation indicates that larger synaptic events transmit more information per vesicle.

**Figure 6.**
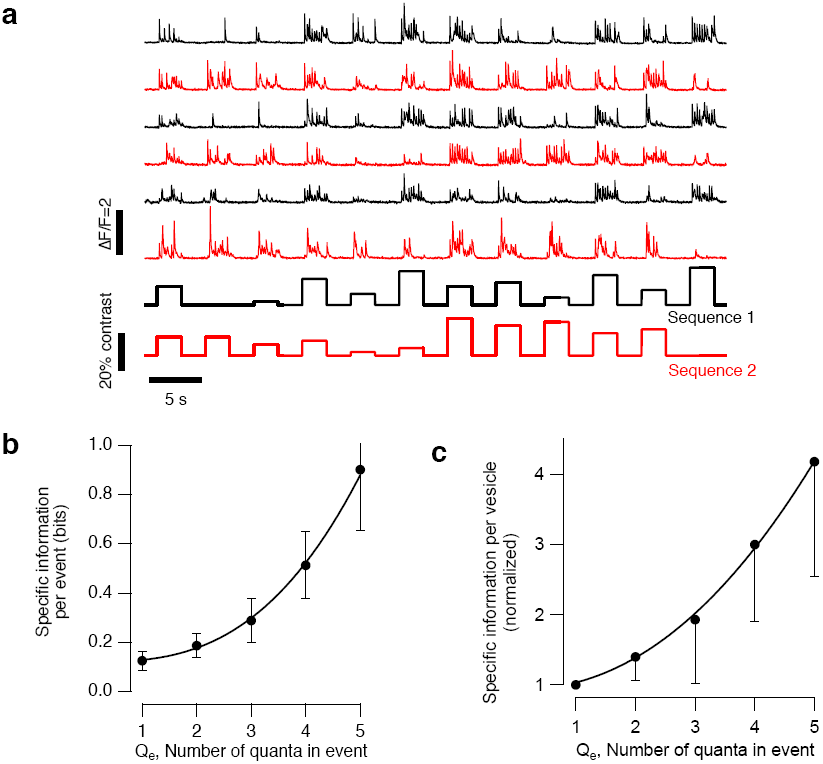
Multivesicular events increased the efficiency of the vesicle code. **a)** Stimulus set for estimating mutual information. Stimuli of varying contrasts were selected to span ± 10% of the range around which the contrast sensitivity was highest in steps of 2%. Note variations in the frequency and amplitude of events at different contrast levels. Stimuli were applied in two different sequences which were repeated alternately. **b)** Specific information per event (i, bits) as a function of Q_e_, the number of vesicles comprising the event. The curve describing the points is a power function of the form i = i_o_ + A.Q_e_^x^, with i_o_ = 0.12 bits, A= 0.008, x = 2.8. Results pooled from 17 synapses. Bars show standard errors. **c)** Specific information per vesicle normalized to the value measured for a uniquantal event (i’). The curve describing the points is a power function of exponent 2.1. Results from the same 17 synapses in **b**.

To explore this idea more directly we divided the amount of information in an event by the number of vesicles it contained to provide a measurement of “vesicle efficiency” before averaging across synapses to quantify the trend. The change in this quantity is plotted in Fig. 6c after normalizing to the value measured in the same synapse for one vesicle. This analysis confirmed that multiquantal events transmit more information *per vesicle*. For instance, events composed of five quanta carried, on average, four times as much information per vesicle as uniquantal events (Fig. 6c). The amplitude code at ribbon synapses counteracts the reduction in the efficiency of information transmission that accompanies the conversion from analogue to digital signaling^15^.

## Discussion

Achieving single-vesicle resolution with iGluSnFR has provided the opportunity to investigate the synaptic code by counting vesicles released from individual active zones, much as electrophysiology has been used to investigate the spike code of neurons^2^. This *in vivo* approach demonstrates that the coordinated release of two or more vesicles is a fundamental aspect of the strategy by which ribbon synapses recode an analogue signal for transmission across the synapse. CMVR discretizes the analogue signal into about eleven output values, with larger modulations of intensity increasing the average quantal content of synaptic events (Fig. 4b and c). As a result, vesicles are used to generate a hybrid code that consists of changes in both the rate and amplitude of events (Figs. 3-5). The amplitude code complements the rate code by increasing the temporal accuracy (Fig. 5), efficiency (Fig. 6) and operating range (Fig. 3d) of synapses transmitting the visual signal to RGCs.

### An amplitude code at ribbon synapses

In essence, CMVR provides a mechanism by which the excitatory effects of glutamatergic vesicles can be summed. Normally, this process occurs on the dendrites of the post-synaptic neuron, with signals from different vesicles adding up on the time scales of ∼5-20 ms over which synaptic potentials decay^35^. An increase in stimulus strength increases the rate of vesicle release and, therefore, the chances of summation. The key distinctions of CMVR are vesicles are summed presynaptically and effectively instantaneous. It will therefore be important to understand how much more effectively single multiquantal events trigger spikes in ganglion cells compared to transient increases in the rate of release of single vesicles. Reliable experimental answers to this question will be difficult to obtain using electrophysiology because a patch pipette records all the current from the compartment it is contacting, making it difficult to distinguish true CMVR from the coincidental arrival of vesicles released from different synapses. A very direct and reliable assessment of the postsynaptic effects of CNVR has, however, been possible in the auditory system of bullfrog where afferents contacting only one hair cell ribbon can be patched. Measuring the currents injected into the afferent by CMVR relative to the currents needed to depolarize the fiber beyond threshold demonstrates that larger glutamatergic events will reliably trigger spikes^20^.

One factor determining how efficiently presynaptic summation of glutamate converts into a postsynaptic current is whether or not post-synaptic receptors glutamate receptors saturate. Saturation does not appear to be an issue at AII amacrine cells postsynaptic to rod-driven bipolar cells^17^ or fibers postsynaptic to hair cells^20^, but it may nonetheless be that there are variations in the efficiency of connections between different cell types. Given the observation that different bipolar cells utilize CMVR to different degrees (Figs. 3 and 4), it may even be that the distribution of sizes of glutamate packets is matched to the properties of receptors on the postsynaptic neuron.

This study demonstrates that the average amplitude of synaptic events is determined by the amplitude of the modulations in light intensity, while previous measurements of CMVR have failed to detect any dependence on the strength of an electrophysiological stimulus^17,20^. One possibility is that the normal calcium-dependence of CMVR was compromised during whole-cell recordings because of wash-out of diffusible intracellular components. For instance, ribbon synapses contain high concentrations of diffusible calcium buffers which act to localize calcium signals around calcium channels and inhibit the release of vesicles docked further away^36^. Wash-out of these buffers through a pipette allows more vesicles to be released by very brief calcium transients and may also disrupt the normal relationship between the calcium current and CMVR. Regenerative calcium signals (“calcium spikes”) within the bipolar cell terminal are also subject to washout^27^ and may be one of the mechanisms which coordinates the release of multiple vesicles. These problems might be overcome by using the perforated patch technique to maintain intracellular molecules while stimulating a bipolar cell or hair cell^36^. Certainly, the relationship between the pre-synaptic calcium signal and the triggering of CMVR requires further investigation.

### Potential mechanisms of CMVR

The cellular mechanisms synchronizing the release of two or more vesicles remain to be discovered. One suggestion is that vesicles attached to the ribbon can fuse to each other to create a multiquantal packet of glutamate which can subsequently fuse with the surface membrane to generate a larger post-synaptic event^16^. This idea is supported by two lines of evidence. Electron microscopy of bipolar cell terminals demonstrates the appearance of large tubular structures on or around the ribbon after a period of stimulation^37,38^ and fluorophore-assisted light inactivation of the ribbon reduces both the rate and amplitude of excitatory post-synaptic potentials^19^. It has been suggested that variable amounts of glutamate might also be released from a single vesicle because of the dynamics of a fusion pore^39^, but this runs counter to evidence that bipolar cells transmit by full collapse of vesicles into the membrane surface^40,41^. Electrophysiological evidence provides stronger support for the idea that ribbon synapses can coordinate the simultaneous fusion of two or more vesicles rather than releasing them all independently^16,17,19^ and this idea is corroborated by the quantization of iGluSnFR events shown in Fig. 1.

The presence of ribbon structures at synapses driven by analogue signals correlates with a second functional specialization - a continuous mode of operation that allows the transmission of sensory information to be maintained over prolonged periods^12,42^(Figs. 4 and 5). The ribbon is thought to support continuous release by capturing vesicles from a mobile pool in the cytoplasm^37^ and then transporting them to the active zone^19^. The maximum rates of continuous release that we observed using iGluSnFR were in the range of 50-100 vesicles s^−1^ per active zone (Fig. 3b), which is in agreement with measurements made using the membrane dye FM1-43^42,43^ and the fluorescent reporter protein sypHy^14^.

### Amplitude and rate coding in relation to the output of the retina

For vision to be useful, information about important stimuli must be transmitted over an appropriately short time window. Multiquantal events were rare and strongly dependent on temporal contrast, so their arrival provided more information about a preceding stimulus than vesicles released individually (Fig. 6). Crucially, CMVR also encoded the timing of a stimulus more precisely: the largest glutamatergic events jittered by just a few milliseconds relative to a stimulus (Fig. 5), which is similar to the spike responses observed in postsynaptic ganglion cells. These spike trains often consist of brief increases in firing rate from longer background periods of silence, making it difficult to describe activity as a time-varying rate. Rather, it has been suggested that “Firing events containing single spikes or bursts of spikes are elicited precisely enough to convey distinct packets of visual information, and hence may constitute the fundamental symbols in the neural code of the retina”^26^. It seems likely that these symbols originate in the amplitude code of bipolar cell synapses.

To understand why a coding strategy based on amplitude might have arisen, it is useful to think about the temporal requirements of a simple rate code. If a Poisson synapse releasing all vesicles independently encodes an event by changing the rate of vesicle release from R to kR, the signal-to-noise ratio achieved over an observation time Dt will be

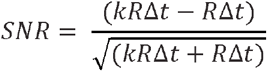

(Online Methods, Equation 4). If one considers the range of contrasts around C_1/2_ ± 10%, the rate of vesicle release was modulated by a factor k of 1.8 (Q_e_ x E_c_; Fig. 3c), from a basal rate R of no more than 20 vesicles s^−1^. To detect such a change with a SNR of 4 (i.e good reliability) would require an observation time of ∼3.5 ms. In comparison, within an OFF synapse, any release event with amplitude greater than four quanta would *immediately* signal an increase in contrast beyond 20% (Fig. 4c). This simple comparison illustrates one of the potential advantages of recoding an analogue signal using symbols varying in amplitude rather than as rates of digital events: an unexpected symbol immediately imparts new information, while a stochastic rate code composed of a single event must be observed over a time window sufficiently long to establish a significant change relative to the noise. Further analysis of the statistics of CMVR will likely shed light on the properties of the vesicle code transmitting visual information.

## Acknowledgments

Many thanks to Jamie Johnston for discussions and to Hazel Smulders and Natasha Bashford for looking after our zebrafish. Funding: Wellcome Trust and BBSRC. Author contributions: This was a team project. S-HS performed molecular biology, fish transgenesis and initial functional analysis; JM-D carried out experiments and analysis and helped prepare the manuscript and supplemental information; LD performed molecular biology, and carried out experiments and analysis; BJ wrote software, conceived and designed information theory experiments and estimates of the transmitter-triggered average, helped prepare the manuscript and supplemental information; LL conceived the project, designed experiments, wrote software, analyzed results and prepared the manuscript.

## Online Methods

### Zebrafish husbandry

Fish were raised and maintained under standard conditions on a 14 h light/10 h dark cycle^14^. To aid imaging, fish were heterozygous or homozygous for the casper mutation which results in hypopigmentation and they were additionally treated with1-phenyl-2-thiourea (200 µM final concentration; Sigma) from 10 hours post fertilization (hpf) to reduce pigmentation further. All animal procedures were performed in accordance with the Animal Act 1986 and the UK Home Office guidelines and with the approval of the University of Sussex local ethical committee.

### Molecular Biology

The zebrafish ribeye a (ctbp2) promoter was used to drive expression of iGluSnFR in all neurons with ribbon synapses (17). Tg(–1.8ctbp2:Gal4VP16_BH) fish that drive the expression of the transcriptional activator protein Gal4VP16 were generated by co-injection of I-SceI meganuclease and endofree purified plasmid into wild-type zebrafish with a mixed genetic background. A myocardium-specific promoter that drives the expression of mCherry protein was additionally cloned into the plasmid to allow for phenotypical screening of founder fish. Tg(10xUAS:iGluSnFR_MH) fish driving the expression of the glutamate sensor iGluSnFR^22^ under the regulatory control of the 10 x UAS enhancer elements were generated by co-injection of purified plasmid and tol2 transposase RNA into offspring of AB wildtype fish outcrossed to casper wildtype fish. The sequences for the myocardium-specific promoter driving the expression of enhanced green fluorescent protein (mossy heart) were added to the plasmid to facilitate the screening process. Crossing the stable lines of Tg(– 1.8ctbp2:Gal4VP16_BH) with Tg(10xUAS:iGluSnFR_MH) results in the targeted expression of the transcriptional activator protein Gal4VP16 that subsequently binds to the UAS elements and switches on the expression of the sensor.

Plasmids for the transgenesis protocol were generated via the Gibson Assembly Cloning method. The table shows the plasmid and primer information, the underlined nucleotide sequences show the part of the primer that anneals to the template during the polymerase chain reaction.

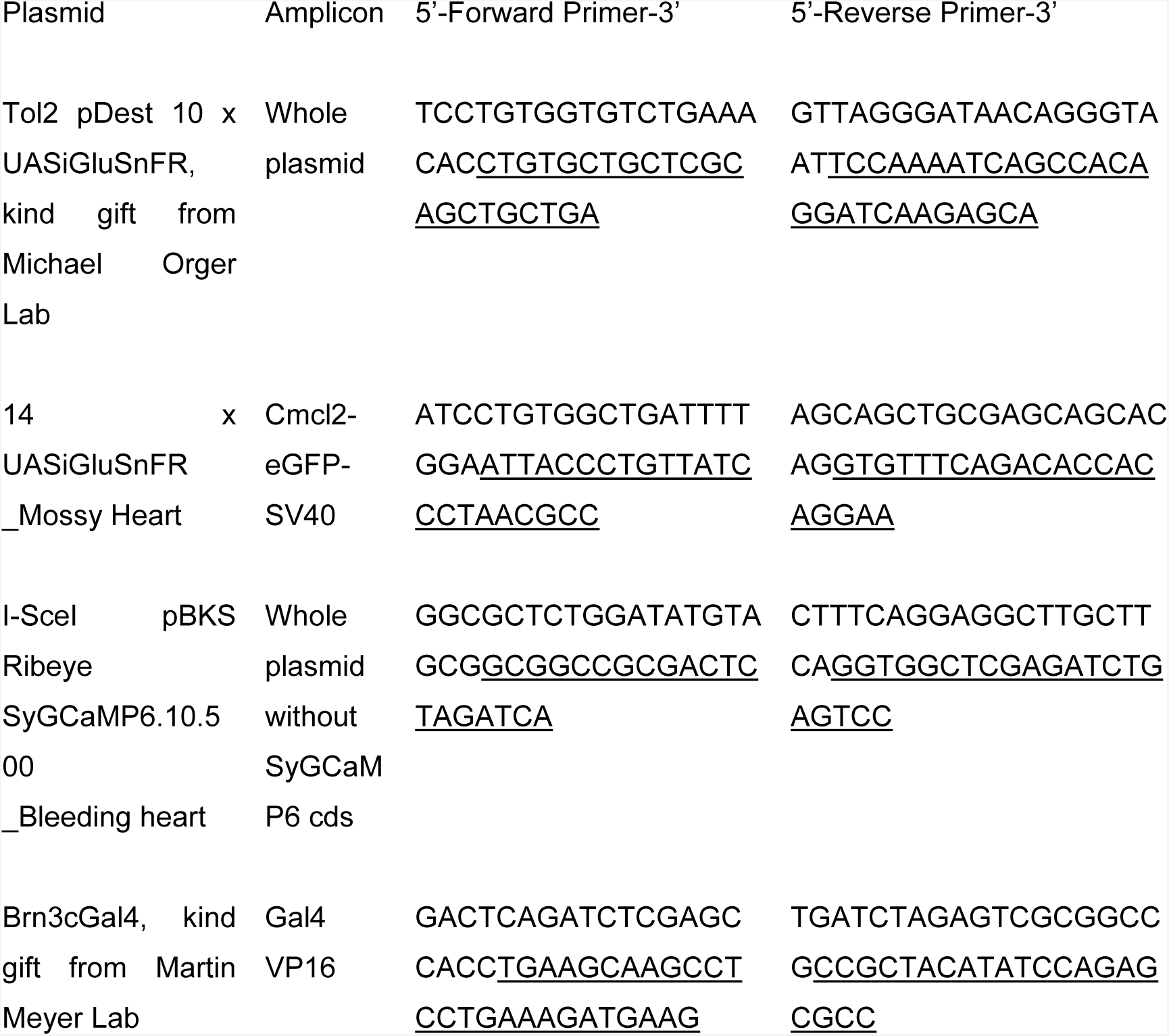

### Multiphoton Imaging *In Vivo*

Zebrafish larvae (7–9 days post-fertilization) were immobilized in 3 % low melting point agarose (Biogene) in E2 medium on a glass coverslip (0 thickness) and mounted in a chamber where they were superfused with E2, as described previously^14^. Imaging of bipolar cell terminals was carried out using a two-photon microscope (Scientifica) equipped with a mode-locked titanium-sapphire laser (Chameleon, Coherent) tuned to 915 nm and an Olympus XLUMPlanFI 20x water immersion objective (NA 0.95). To prevent eye movements, the ocular muscles were paralyzed by injection of 1 nL of α-bungarotoxin (2 mg/mL) behind the eye. The signal-to-noise ratio for imaging was optimized by collecting photons through both the objective and a sub-stage oil condenser (Olympus, NA 1.4). Emission was filtered through GFP filters (HQ 535/50, Chroma Technology) before detection with GaAsP photomultipliers (H7422P-40, Hamamatsu). The signal from each detector passed through a current-to-voltage converter and then the two signals were added by a summing amplifier before digitization. Scanning and image acquisition were controlled under ScanImage v.3.6 software^44^. Images were typically acquired at 10 Hz (128 × 100 pixels per frame, 1 ms per line) while linescans were acquired at 1 kHz.

Full-field light stimuli were generated by an amber LED (lmax = 590 nm, Thorlabs), filtered through a 590/10 nm BP filter (Thorlabs), and delivered through a light guide placed close to the eye of the fish. Stimuli were normally delivered as modulations around a mean intensity of ∼165 nW/mm^2^ and intensity was modulated through Igor Pro software (Wavemetrics) running mafPC (courtesy of M. A. Xu-Friedman) through an ITC-18 DAC interface (HEKA). The multiphoton microscope was synchronized to visual stimulation.

### Extraction of glutamatergic events

An analysis package was created within Igor Pro (Wavemetrics) to detect and quantize glutamatergic events from two-photon line-scans across the terminals of bipolar cells expressing iGluSnFR. The analysis comprised of six major steps which are described in detail in the Supplementary Information:

1. Separation of ROIs by spatial decomposition
2. Time series extraction by weighted averaging
3. Baseline correction and calculation of ΔF/F
4. Identification of events by Wiener deconvolution
5. Extraction of events
6. Amplitude clustering to create a time series of quantized events

### Calculation of temporal jitter

In order to compute the temporal jitter of the glutamatergic events, we first calculated the vector strength:

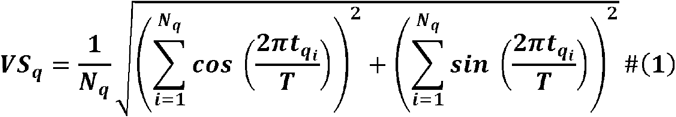

where t_qi_ is the time of the i^th^ q-quantal event, T is the stimulus period, and N_q_ is the total number of events of composed of q-quanta. The temporal jitter can then be computed by:

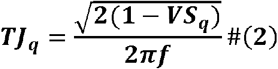

where f is the stimulus frequency.

### The signal-to-noise ratio associated with a change in the rate of a Poisson process

Imagine the mean rate of a Poisson process, R, changes by a factor k. The signal, S, generated by comparing two observation times Dt will be the change in the mean number of events counted in each period (kR Dt - R Dt), and the variance in that signal will be the sum of the number of events counted in each period (kR Dt + R Dt). Defining the SNR in the same way as the discriminability (d’) used in signal detection theory^45^, we have

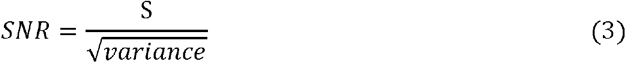

yields

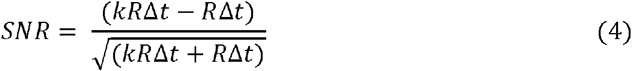

which can be rearranged to obtain the time period Dt required to obtain a given SNR, as

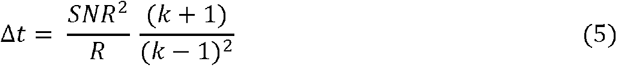

### Calculation of the Transmitter Triggered Average (TTA)

The TTA for an event containing a specific number of quanta (s_q_) was calculated by averaging the stimuli that preceded each release event of that type:

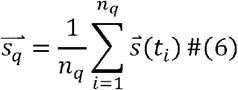

where q is the quantal event type, n_q_ is the number of n-quantal events within the recording, and s(t) is the stimulus window ending at time t. Thus, for instance, a synapse that releases up to 11 vesicles would in principle produce 11 filters, one for each quantal event type. Two different stimuli were used to drive the cell for the TTA analysis. We used Gaussian White Noise (GWN) in which intensities were drawn from a Gaussian distribution spanning the LED drivers input range and updated at a specific stimulus frame rate (varied between 10 and 30 Hz). The GWN was then discretized into eight equally spaced bins to produce a stimulus that approximates a Gaussian distribution while driving synaptic responses effectively.

### Calculations based on information theory

To quantify the amount of information about a visual stimulus that is contained within the sequence of release events from an active zone we first needed to convert bipolar cell outputs into a probabilistic framework from which we could evaluate the specific information (I_2_), a metric that quantifies how much information about one random variable is conveyed by the observation a specific symbol of another random variable^21^. The time series of quantal events was converted into a probability distribution by dividing into time bins of 20 ms, such that each time bin contained either zero events or one events of an integer amplitude. We then counted the number of bins containing events of amplitude 1, or 2, or 3 etc. By dividing the number of bins of each type by the total number of bins for each different stimulus, we get the conditional distribution of Q given S, *p*(*Q*|*S*), where Q is the random variable representing the *quanta/bin* and S is the random variable representing the *stimulus contrasts* presented throughout the course of the experiment. We then compute the joint probability distribution by the chain rule for probability (given the experimentally defined uniform distribution of stimuli S):

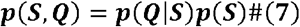

In order to convert this distribution into the conditional distribution of S given Q, we use the definition of the conditional distribution:

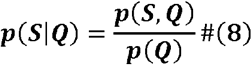

From these distributions we can now compute the specific information as the difference between the entropy of the stimulus S minus the conditional entropy of the stimulus given the observed symbol in the response q:

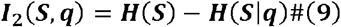

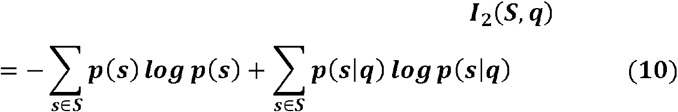

representing the amount of information observing each quantal event type q ▫ Q carries about the stimulus distribution S.

